# Parent of origin gene expression in a founder population identifies two new imprinted genes at known imprinted regions

**DOI:** 10.1101/344457

**Authors:** Sahar V. Mozaffari, Michelle M. Stein, Kevin M. Magnaye, Dan L. Nicolae, Carole Ober

**Author notes:** Corresponding author (SVM).

## Abstract

Genomic imprinting is the phenomena that leads to silencing of one copy of a gene inherited from a specific parent. Mutations in imprinted regions have been involved in diseases showing parent of origin effects. Identifying genes with evidence of parent of origin expression patterns in family studies allows the detection of more subtle imprinting. Here, we use allele specific expression in lymphoblastoid cell lines from 306 Hutterites related in a single pedigree to provide formal evidence for parent of origin effects. We take advantage of phased genotype data to assign parent of origin to RNA-seq reads in individuals with gene expression data. Our approach identified known imprinted genes, two putative novel imprinted genes, and 14 genes with asymmetrical parent of origin gene expression. We used gene expression in peripheral blood leukocytes (PBL) to validate our findings, and then confirmed imprinting control regions (ICRs) using DNA methylation levels in the PBLs.

**Author Summary:** Large scale gene expression studies have identified known and novel imprinted genes through allele specific expression without knowing the parental origins of each allele. Here, we take advantage of phased genotype data to assign parent of origin to RNA-seq reads in 306 individuals with gene expression data. We identified known imprinted genes as well as two novel imprinted genes in lymphoblastoid cell line gene expression. We used gene expression in PBLs to validate our findings, and DNA methylation levels in PBLs to confirm previously characterized imprinting control regions that could regulate these imprinted genes.

## Introduction

Imprinted genes have one allele silenced in a parent of origin specific manner. In humans, approximately 105 imprinted loci have been identified, many of which play important roles in development and growth[1-3]. Dysregulation of imprinted genes or regions can cause diseases that show parent of origin effects, such as Prader-Willi or Angelman syndrome, among others [2]. Imprinted regions have also been associated with complex traits, such as height and age of menarche [4,5], as well as common diseases such as obesity and some cancers [2]. More than 80% of imprinted genes in humans are clustered in genomic regions that contain both maternally and paternally expressed genes, as well as genes that encode non-coding RNAs[2,6]. Parent-specific expression of the genes within a cluster are maintained by complex epigenetic mechanisms at cis-acting imprinting control regions (ICRs) [3], which show parent of origin specific DNA methylation patterns and chromatin modifications[7].

Using RNA-seq and allele specific expression (ASE) we can map genes to parental haplotypes and identify those that are expressed when inherited from only the father or only from the mother, a hallmark feature of imprinted loci. Parent of origin effects and imprinted genes have been most elegantly studied in mice, where two inbred strains are bred reciprocally to identify parent of origin effects on gene expression in progeny that have the same genotypes but different patterns of inheritance [8]. Additionally, uniparental inheritance of imprinted regions in mice were associated with abnormal developmental phenotypes [9] before it was shown that imprinting defects are associated with human disease[10,11]. One approach to identifying imprinted loci in humans has been to test for parent of origin effects on gene expression and phenotypes in pedigrees [4,12]. For example, Garg et al. used gene expression in LCLs from HapMap trios to identify 30 imprinting eQTLs with parent of origin specific effects on expression [13]. A study from the GTEx Consortium used RNA-seq data and allele specific expression to identify allelic imbalance in 45 different tissues. By considering genes with monoallelic expression that was evenly distributed to both the reference and alternate alleles across individuals as evidence for imprinting, they identified 42 imprinted genes, both known and novel, and used family studies to confirm imprinting of 5 novel imprinted genes [14]. Santoni et al. identified nine novel imprinted genes using single-cell allele-specific gene expression and identifying genes with mono-allelic expression in fibroblasts from 3 unrelated individuals and probands of 2 family trios, and then used the trios to confirm parent of origin of the alleles [15].

Here, we perform a parent of origin ASE study in a large pedigree to characterize parent of origin specific gene expression in the Hutterites, a founder population of European descent, for which we have phased genotype data [16]. We use RNA-seq from lymphoblastoid cell lines (LCLs) to map transcripts to parental haplotypes and identify known and two not previously reported imprinted genes. We validated the two putative imprinted genes by showing the same patterns of parent of origin expression PBLs from different Hutterite individuals, and show DNA methylation signatures of imprinting in the PBLs at these regions.

## Results

### Mapping transcripts to parental haplotypes

For each of 306 individuals, the total number of transcripts at each gene was assigned as maternally inherited, paternally inherited, or unknown parent of origin. The last group included transcripts without heterozygote SNPs or transcripts with SNPs without parent of origin information. Transcripts were assigned to the parentally inherited categories using SNPs in the reads and matching alleles to either the known maternally or paternally inherited alleles. All the genes analyzed had some transcripts of unknown origin (average 97.8%, range 8.3-100%). For each gene we assigned parental origin to an average of 1.8% of transcripts (range: 0-34.7%), and for each individual we assigned parental origin to an average of 1.4% of transcripts (range: 0-1.7%). On average, about 40 SNPs per gene were used to assign the transcripts of a gene to parent (range 1-1839 SNPs).

**Table 1.** Summary Statistics for Parental Origin of Transcripts.

|  | Mean | Standard Deviation | Range |
| --- | --- | --- | --- |
| Proportion of transcripts from each gene assigned to transcripts of unknown origin | 0.978 | 0.031 | (0.083, 1) |
| Proportion of transcripts from each gene assigned to parental origin | 0.018 | 0.019 | (0, 0.347) |
| Proportion of transcripts for each individual assigned to parental origin | 0.014 | 0.0015 | (0, 0.017) |

After quality control (see Methods), transcripts in 15,889 genes were detected as expressed in 306 individuals. Some transcripts for 14,791 of those genes could be assigned to a parent. Of these, 75 genes were only expressed on the paternally-inherited allele in at least one individual and not on the maternally inherited allele in any individuals. Similarly, 64 genes were only expressed on the maternally-inherited allele in at least one individual and not on the paternally inherited allele in any individuals (S1 Table).

### Imprinted Genes in Lymphoblastoid Cell Lines (LCLs)

Among the 139 genes with only paternally inherited expression or only maternally inherited expression, there are three known imprinted genes (*CDKN1C*, *NDN*, *SNRPN*) and one previously predicted to be imprinted (*IFITM1*) [17]. *CDKN1C* showed patterns opposite of what has been reported [18,19], which could be due to the small sample (only three individuals showed expression from one parent) or to the different cell types used here (LCLs) and in previous studies (developing brain and embryonal tumors for *CDKN1C*).

We expect some imprinted genes to have ‘leaky’ expression, such that there is some expression from the parental chromosome that is mostly silenced. To detect these genes, we used a binomial test to find patterns of gene expression asymmetry by parental transcript levels. This analysis identified 28 genes with an FDR <5% (Table 2). The 11 genes that showed the most asymmetry are known imprinted genes: *ZDBF2*, *PEG10*, *SNHG14*, *NHP2L1*, *L3MBTL1*, *ZNF331*, *LPAR6*, *FAM50B*, *KCNQ1*, *NAP1L5*, and *IGF1R*. Parent of origin expression for *ZDBF2* and *KCNQ1* are shown in **Fig 1A** and **1B**, respectively. We identified two additional genes that showed asymmetry in parental expression from mostly one parent (*PXDC1, PWAR6*), which we consider potentially new imprinted genes. The remaining fourteen genes showed significant patterns of asymmetry but had expression from both maternal and paternal chromosomes. These genes are likely not imprinted but could have asymmetry in expression due to an expression quantitative trait loci (eQTL).

**Figure 1.**
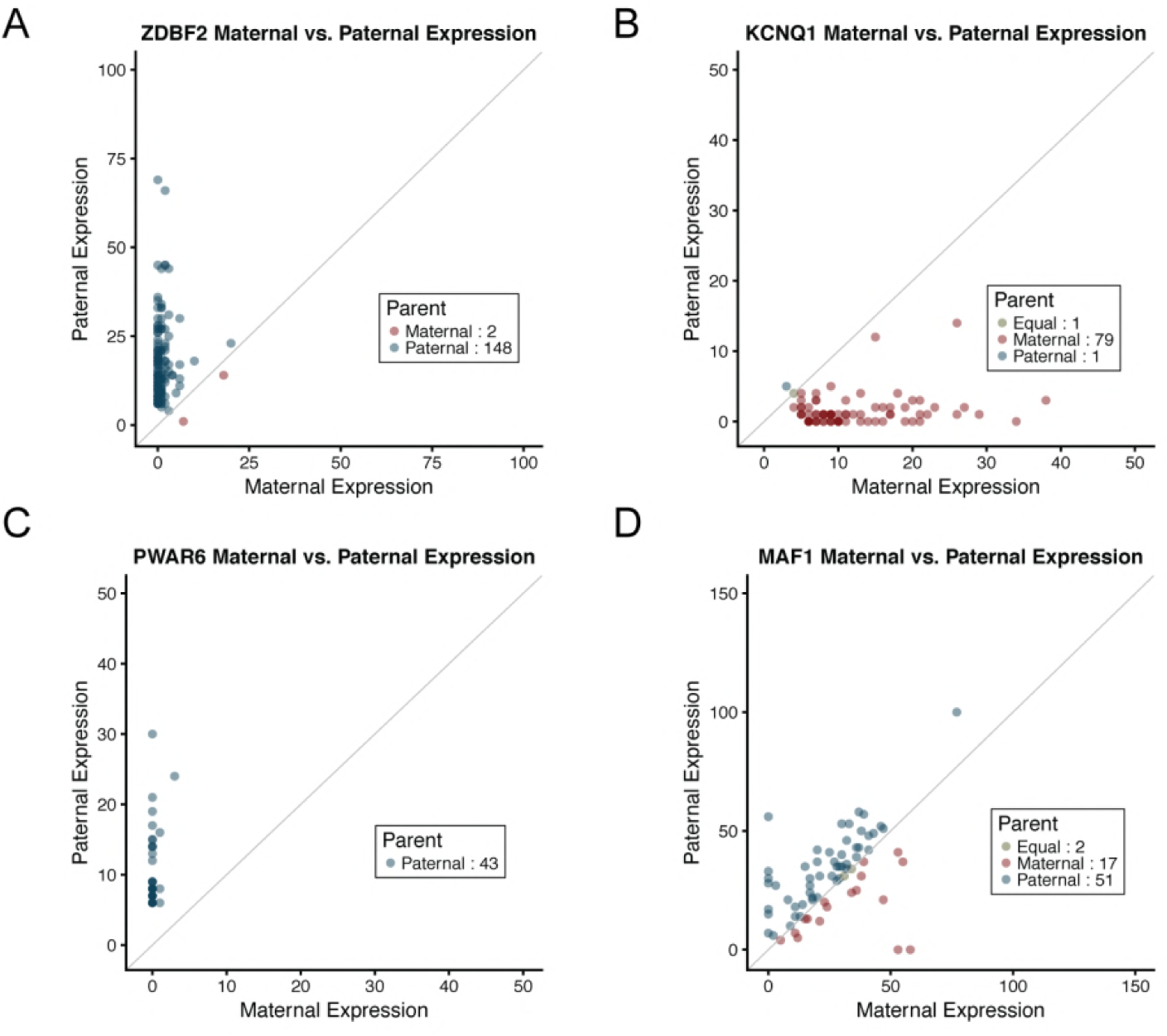
Plot of maternal (x-axis) and paternal (y-axis) gene expression for four genes. (**A)** maternally imprinted gene *ZDBF2* (paternally expressed), (**B)** paternally imprinted gene *KCNQ1* (maternally expressed), (**C)** novel maternally imprinted gene *PWAR6* (paternally expressed), (**D)** gene with asymmetry in parental expression *MAF1*. Each point represents one individual. Numbers in the legend represent the number of individuals with equal maternal and paternal expression, more maternal expression, or more paternal expression.

**Table 2.**
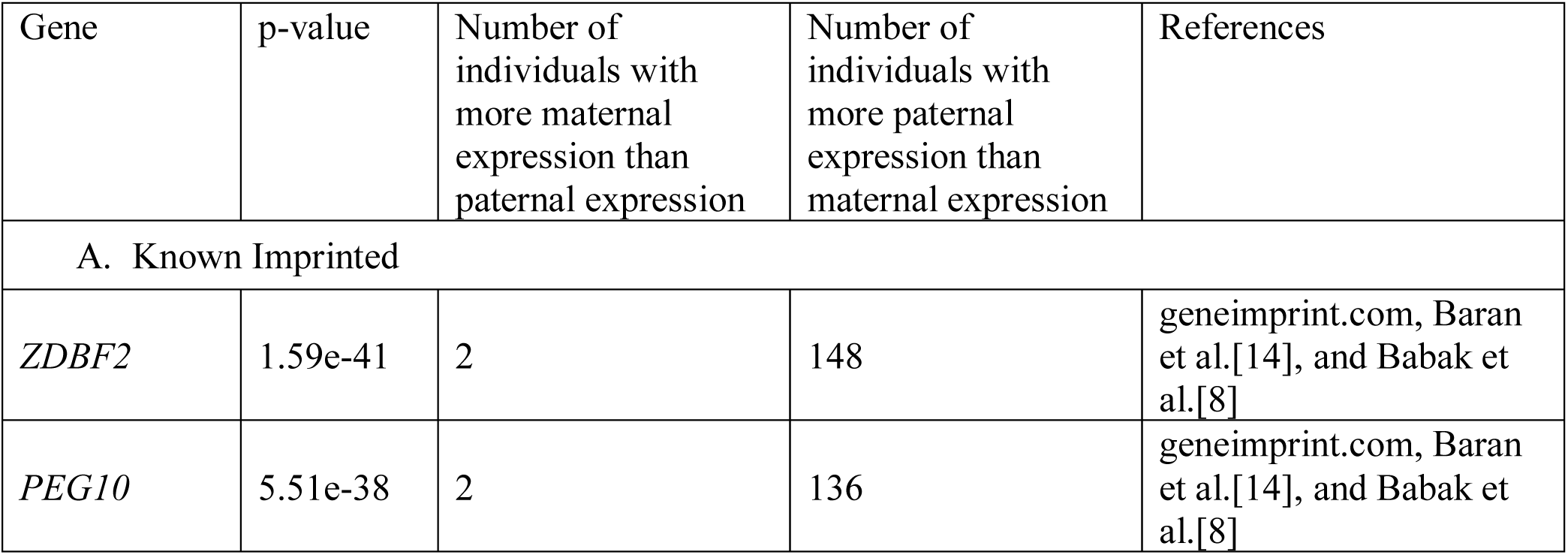
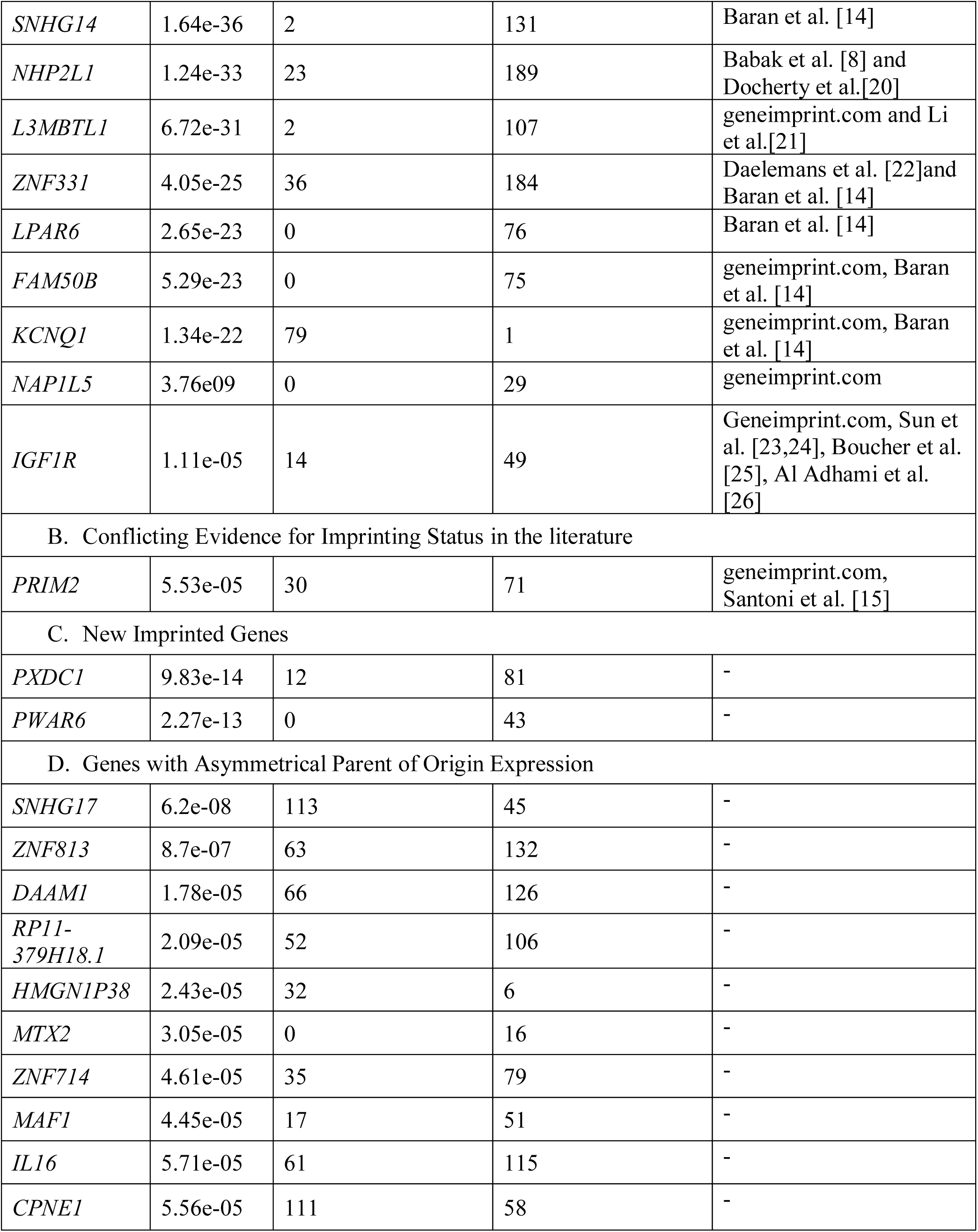
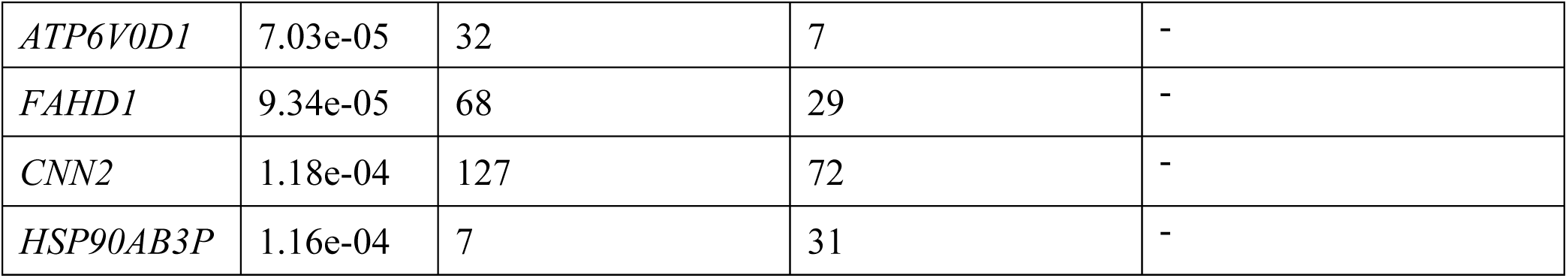
Results for Gene with Parent of Origin Expression Asymmetry. Genes listed by category of imprinting status: (A) Known Imprinted, (B) Conflicting Evidence for Imprinted Status, (C) New Imprinted Genes, (D) Genes with Asymmetrical Parent of Origin Expression. Genes are ordered by significance within each category.

Two genes showed gene expression signatures consistent with imprinting but have not previously been recognized as imprinted genes. The first potentially new imprinted gene is *PXDC1*, which is in the same region and next to (<100kb) a known imprinted gene, *FAM50B*. The second potentially novel imprinted gene is *PWAR6*, or Prader Willi Angelman Region RNA6, a gene encoding a regulatory class of RNA. Although this gene is located within the intron of a known imprinted gene, *SNHG14*, this noncoding RNA has not previously been recognized as having parent of origin specific expression (**Fig 1C**).

The remaining fourteen genes show significant asymmetry using the binomial test but do not have expression from mostly one parental chromosome. One of these genes, *SNHG17*, is a noncoding RNA. Another gene with parent of origin asymmetry, *ZNF813*, is next to a known imprinted gene, *ZNF331*. The remaining genes with asymmetrical parent origin expression have expression from both parental chromosomes, unlike imprinted genes. These genes include *DAAM1*, which is involved in cytoskeleton, specifically filopodia formation [27,28], and has a suggested role for cytoskeleton organization during Mammalian testis morphogenesis and gamete progression [29]; *RP11-379H18.1*, a noncoding RNA gene; *HMGN1P38* [30]; *MTX2*, a nuclear gene that interacts with mitochondrial membrane protein metaxin 1 and is involved in mitochondrial protein import and metabolism of proteins in mice; *MAF1*, a negative regulator of RNA polymerase 2; *ZNF714*, *CPNE1*, *IL16*, *ATP6V0D1*, *FAHD1*, *HSP90AB3P*, and *CNN2* are the remaining genes that show parent of origin asymmetry but not with a pattern consistent with imprinting (**S1 Figure**).

### Validation of Imprinted Genes in PBLs

Using the same methods described above, we assigned parent of origin to transcripts in PBLs from 99 Hutterite individuals not included in the LCL studies. Maternal and paternal expression in PBLs for all 28 genes identified in LCLs showed similar trends of asymmetry as in LCLs (**Fig 2**).

**Figure 2.**
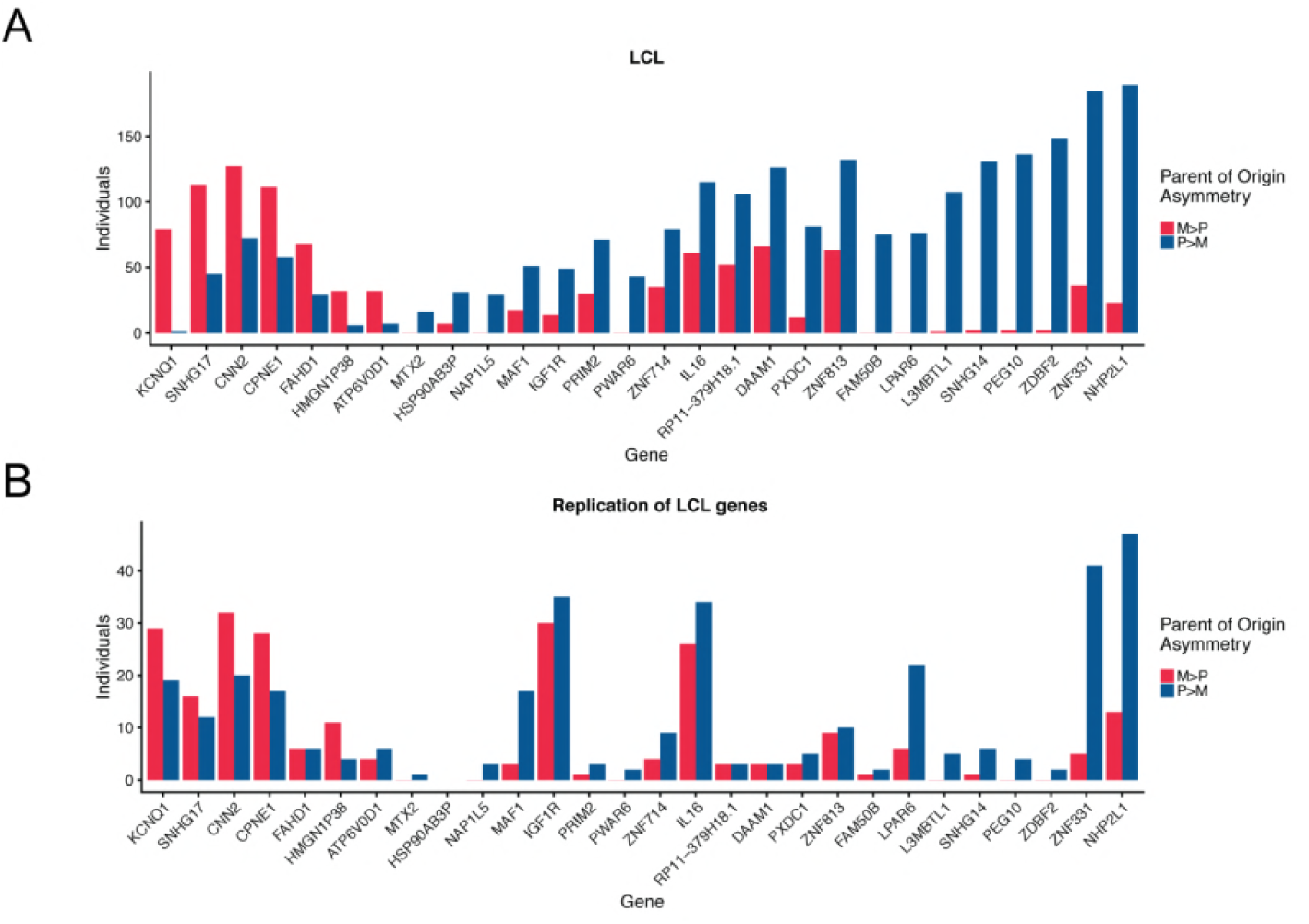
Histogram showing the number of individuals with more maternal expression (M>P) or more paternal expression (P>M) for the 28 genes showing parent of origin asymmetry in (**A**) LCLs and (**B**) PBLs. Genes are ordered by the magnitude of the difference in the number of individuals with more maternal expression than paternal expression in LCLs.

### Methylation at Imprinting Control Regions

One of the mechanisms underlying parent of origin effects on expression at imprinted loci is differential methylation at cis-acting imprinting control regions (ICRs). DNA methylation from the Illumina HumanMethylation 450K array was available in PBLs from the same individuals included in the validation study described above. To determine the expected patterns of methylation at known imprinted loci, we first looked at previously characterized methylated regions at known imprinted regions from Court et al. and Joshi et al. [31,32].

The methylation patterns at the two potentially novel imprinted genes identified in this study, *PXDC1* and *PWAR6*, lie in or near known imprinted regions that contain previously characterized ICRs. These previously characterized ICRs show about 50% methylation (beta value of between 0.25 and 0.75) in our DNA methylation data, which likely reflect methylation at only one parental chromosome in all the cells in the sample. Methylation patterns in PBLs at these two ICRs fall within this hemi-methylation range, further suggesting that these two genes are indeed imprinted (**Fig 3**).

**Figure 3.**
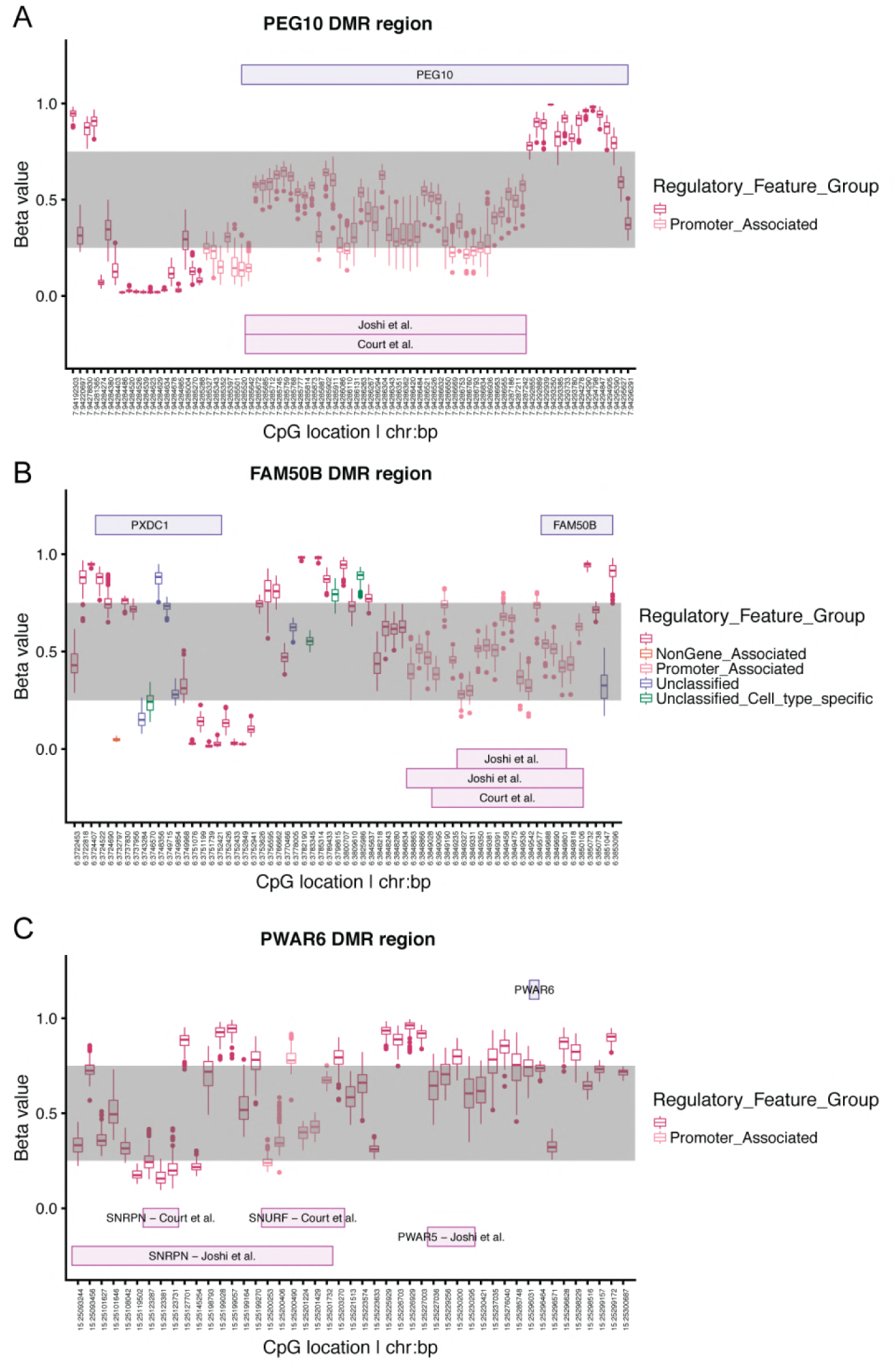
DNA methylation levels near known and novel imprinted genes previously defined by Joshi et al. and Court et al. (**A)** *PEG10*, (**B)** *PXDC1* and *FAM50B*, (**C)** *PWAR6.*

## Discussion

Dysregulation of imprinted genes can have a large impact on mammalian development and has been associated with significant diseases in humans. Studies aimed at identifying imprinted genes at genome-wide levels have used allele specific expression and imbalance to infer parent of origin. Here we used a large pedigree with assigned parent of origin alleles to map transcripts to chromosomes with known parent of origin and identify imprinted genes.

Using this approach, we found genes with expression primarily from either the maternal or paternal haplotype. Because gene silencing at imprinted loci may be incomplete, we used a binomial test on parent of origin gene expression and identified 11 known imprinted genes and two potentially novel imprinted genes. Both of these novel genes, *PWAR6* and *PXDC1*, lie in known imprinted regions but have not themselves been characterized as imprinted. The remaining genes that have significant parent of origin asymmetry in gene expression do not show clear imprinting expression patterns. To validate these findings, we mapped gene expression in PBLs from Hutterite individuals not included in the LCL study. The same genes showed similar patterns of asymmetry in these different cell sources (transformed B cells and peripheral blood leukocytes) from different individuals.

We also characterized methylation patterns near genes showing asymmetry. Using results from studies that had previously characterized ICRs in patients with uniparental disomy at many imprinted regions [31,32], we estimated regions for defining hemi-methylation near the genes identified in our study. Using this approach, we were able to provide additional supportive data for the two potentially new imprinted genes to be true imprinted genes regulated by previously characterized ICRs.

Although our study is the largest pedigree-based study to date to search genome-wide for imprinted genes, it has limitations. First, we are able to determine the parent of origin for a many transcripts in the Hutterites but we could not assign every RNA sequencing read to a parent due to lack of heterozygous sites or missing parent of origin information for alleles. Second, we conducted these studies in lymphoblastoid cell lines, and therefore could only study genes imprinted in this cell type and would miss the many imprinted genes that are tissue-specific and/or developmentally regulated[33]. Third, while we can verify previously characterized ICRs, our study is not designed to identify novel ICRs because DNA methylation values from an array cannot be assigned to parental haplotype. Lastly, although we characterized the gene expression and methylation patterns for two potentially novel imprinted genes, replication of these genes in a different population and in different tissues, and functional characterization of these genes are required to confirm their status as imprinted genes. Similarly, some of the other genes with parent of origin asymmetry in the blood cells examined in this study may show more clear-cut evidence for imprinting in other tissues or at specific periods of development.

In summary, we have identified two new imprinted genes using gene expression from a founder population. The genes with asymmetrical parental expression had similar patterns of asymmetry in a different source of blood cells and in different individuals, and we were able to replicate the methylation patterns in known ICRs near the known and novel imprinted genes in this study. Our method and study population allowed us to map reads to parental haplotypes and uncovered *PWAR6* and *PXDC1* as new imprinted genes that could potentially impact disease risk and development.

## Methods

### Genotypes

Hutterite individuals (n=1,653) were genotyped using one of three Affymetrix genotype arrays, as previously described [16], of which 121 underwent whole genome sequencing by Complete Genomics, Inc (CGI) (n=98) or Illumina whole genome sequencing (n=27). A total of 10,235,233 variants present in the sequenced individuals were imputed and phased to the remaining 1,532 genotyped individuals using PRIMAL [16]. Parent of origin was assigned to 89.85% of the alleles with call rate 81.6842% after QC. For this study, we included individuals with genotyped parents in the primary analyses in LCLs. Written consents for these studies were obtained from the adult participants and parents of children under 18; written assents were obtained from all children. This study was approved by the University of Chicago Institutional Review Board.

### RNA-seq in Lymphoblastoid Cell Lines (LCLs)

RNA-seq was performed in LCLs as previously described [34]. For this study, sequencing reads were reprocessed as follows. Reads were trimmed for adaptors using Cutadapt (reads less than 5 bp discarded) then remapped to hg19 using STAR indexed with gencode version 19 gene annotations [35,36]. To remove mapping bias, reads were processed and duplicate reads removed using WASP [37]. We used a custom script modified from WASP to separate reads that overlap maternal alleles or paternal alleles. Reads without informative SNPs (homozygous, or no parent of origin information) were categorized as unknown where the unknown, maternal, and paternal make up the total gene expression. Gene counts were quantified using STAR for each category. VerifyBamID was used to identify sample swaps [38]. Genes mapping to the X and Y chromosome were removed; genes with a CPM log transformed value less than 1 in less than 20 individuals were also removed.

### RNA-seq in Peripheral Blood Leukocytes (PBLs)

RNA-seq was performed in whole blood as previously described [39]. For this study, sequencing reads were reprocessed as described above for the studies in LCLs. For these analyses, we excluded 32 individuals who were also in the LCL study.

### Identifying Imprinted Genes

We used a binomial test to detect asymmetry in parent of origin gene expression. Using the paternally and maternally assigned reads, we generated a binomial Z-score for each individual for each gene (Z_i_) and excluded those where Z_i_=0. For each gene, the number of subjects with Z_i_ >0 can be modeled by a Binomial distribution with probability ½, under the null hypotheses of symmetric expression. For imprinted genes that show patterns of asymmetry, we expect a distribution of Z-scores that are skewed to one direction corresponding to asymmetric expression. Because we are only asking whether there are more individuals with more maternal expression or more paternal expression and not gene expression measures there is no need to model over-dispersion.

### DNA methylation profiling and processing in PBLs

One milliliter of whole blood from 145 Hutterites was drawn into TruCulture (Myriad RBM; Austin, Texas) tubes containing proprietary TruCulture media. DNA was extracted using AllPrep DNA/RNA Mini Kits (Qiagen). DNA samples were bisulfite converted and hybridized to the Illumina HumanMethylation 450K array at the University of Chicago Functional Genomics Center. Samples were processed using default parameters using the R package minfi [40], normalized using SWAN (subset within-array normalization [41]) and quantile normalized similar to previous methylation studies [42]. Probes were removed if: (1) mapped non-uniquely to a bisulfite-converted genome; (2) mapped to sex chromosomes; (3) had a probe detection p-value >0.01 in at least 25% of samples; and (4) contained common SNPs within the probe sequence, as previously described [43]. Principal components analysis (PCA) was used to identify significant technical covariates, and the ComBat function [44] within the R package sva [45] was used to correct for chip effect. Analyses of DNA methylation levels were conducted using beta values, which were converted from M-values using the lumi R package [46].

## Acknowledgements

We thank members of the Ober lab for useful discussions, Joe Urbanski and Lorenzo Pesce for assistance using Beagle, the many members of our field trip teams, and the Hutterites for their continued support of our studies.

## Funding Disclosure on Submission System

This work was supported by NIH grants HL085197 and HD21244; and in part by NIH through resources provided by the Computation Institute and the Biological Sciences Division of the University of Chicago and Argonne National Laboratory, under grant 1S10OD018495-01. S.V.M has been supported by NIH Grant T32 GM007197 and the Ruth L. Kirschstein NRSA Award F31HL134315.

## Supporting Information

**S1 Figure.**
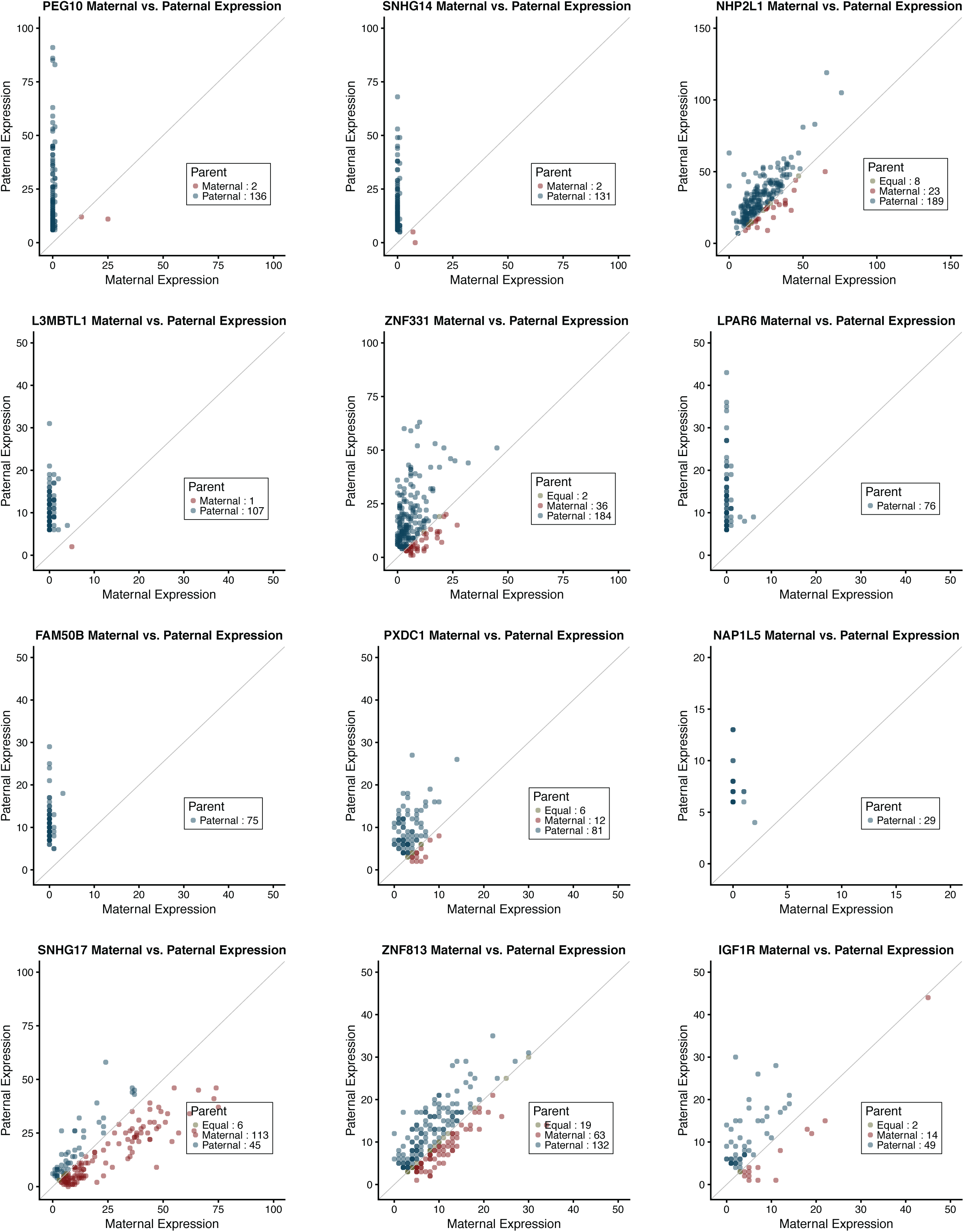
Plots of maternal and paternal expression for remaining genes with parent of origin asymmetry.

**S2 Figure.**
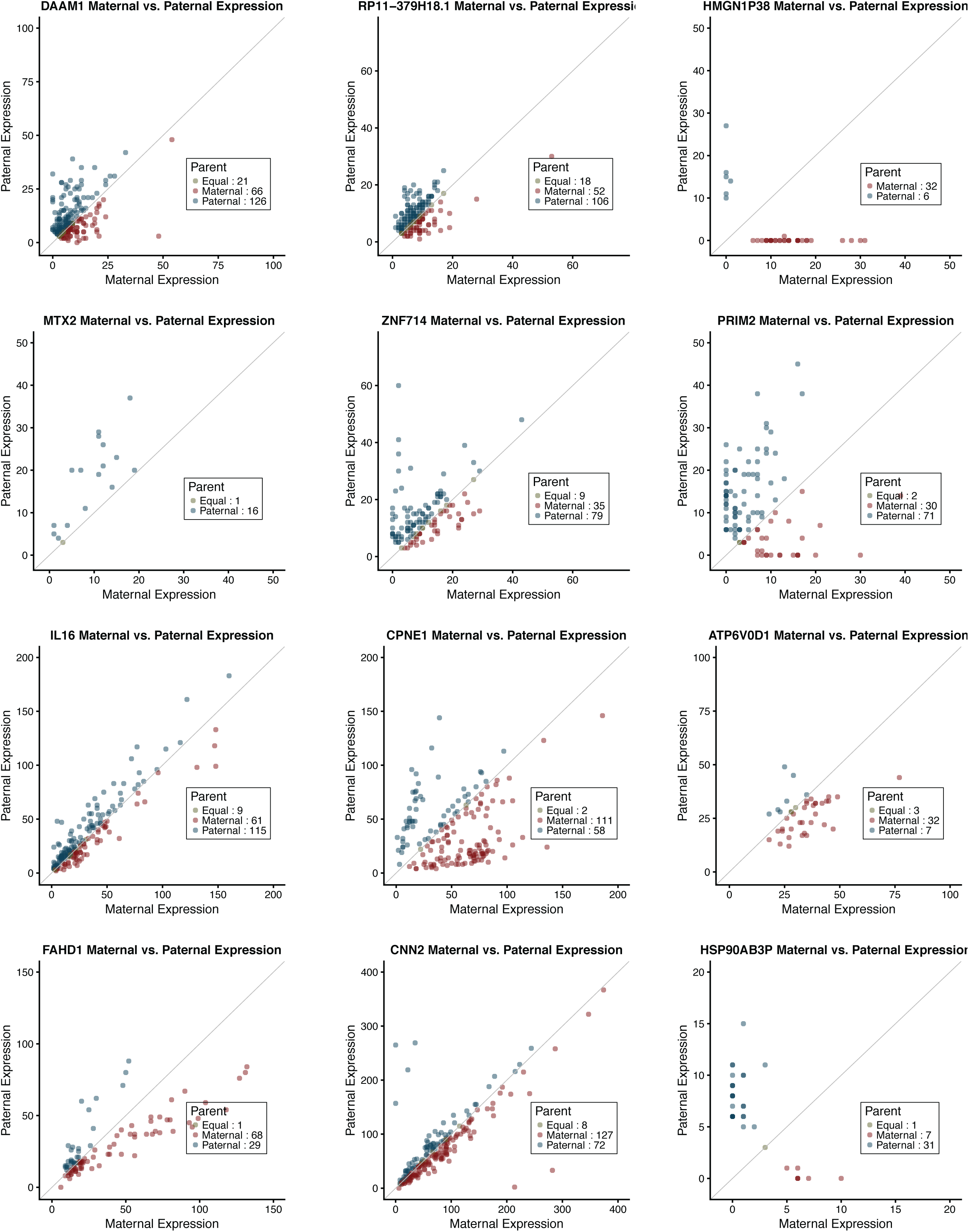
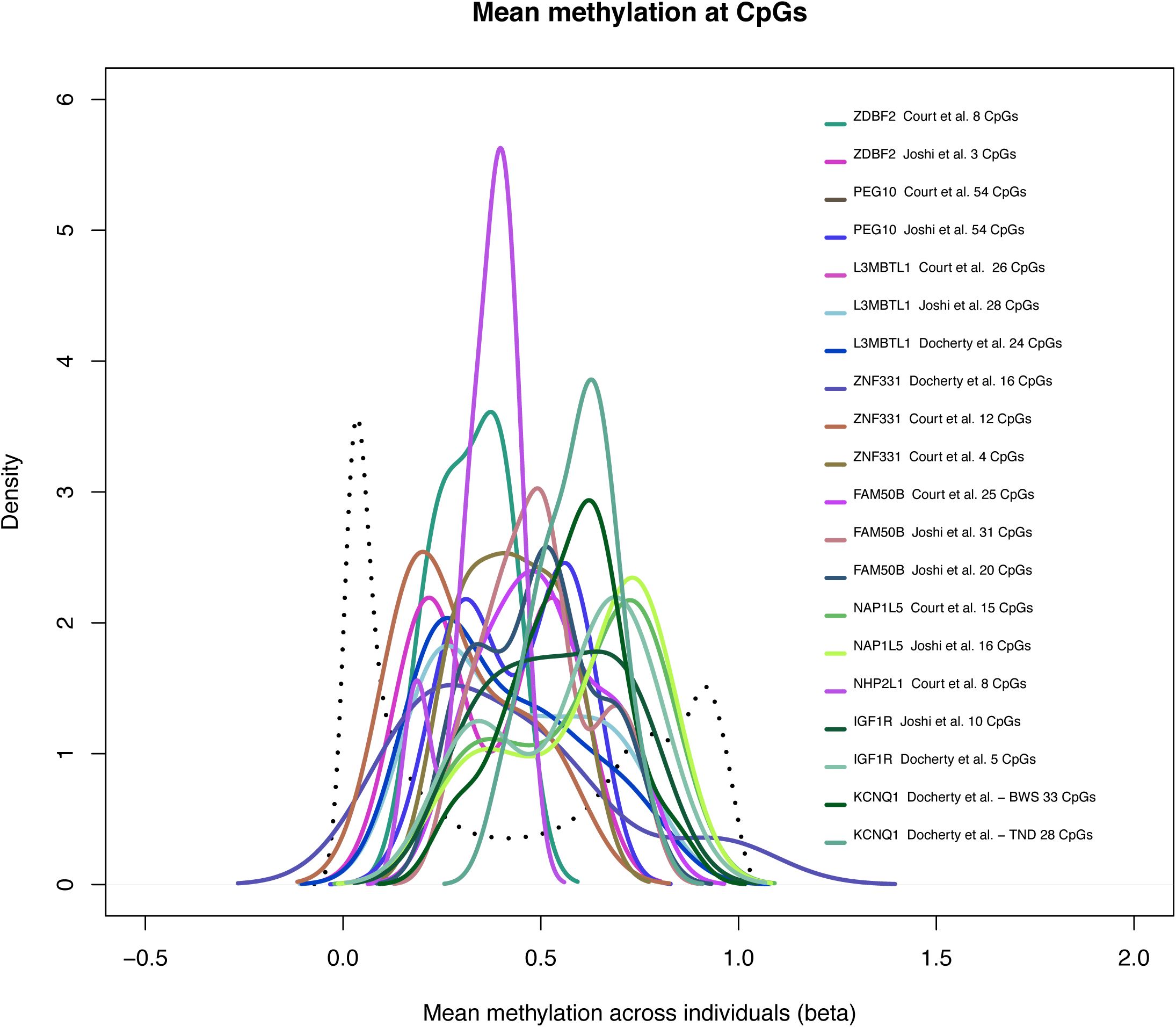
Density plot for Differentially Methylated Regions (DMRs) for imprinted genes from Joshi et al and Court et al.

**Supplementary Table 1.**
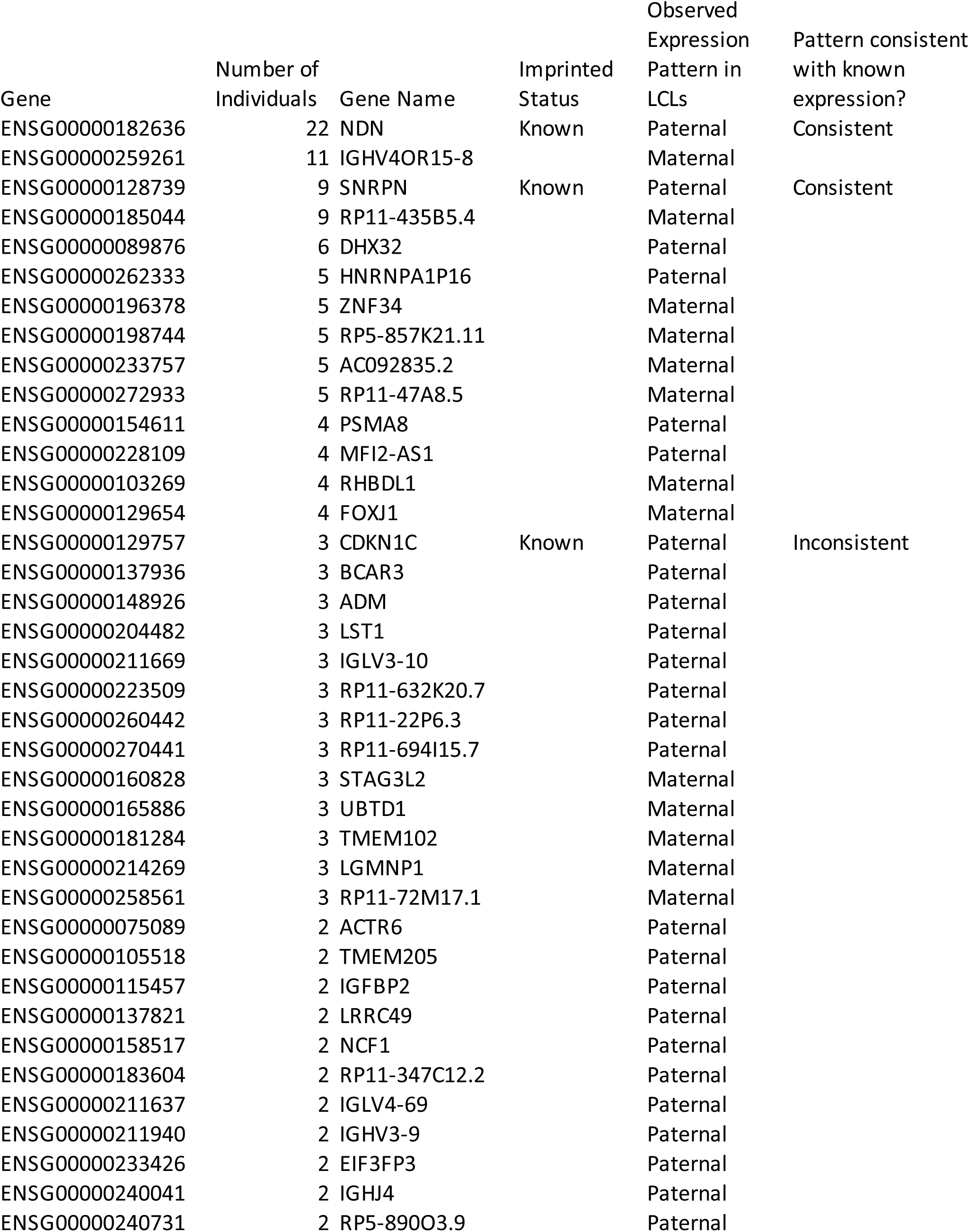
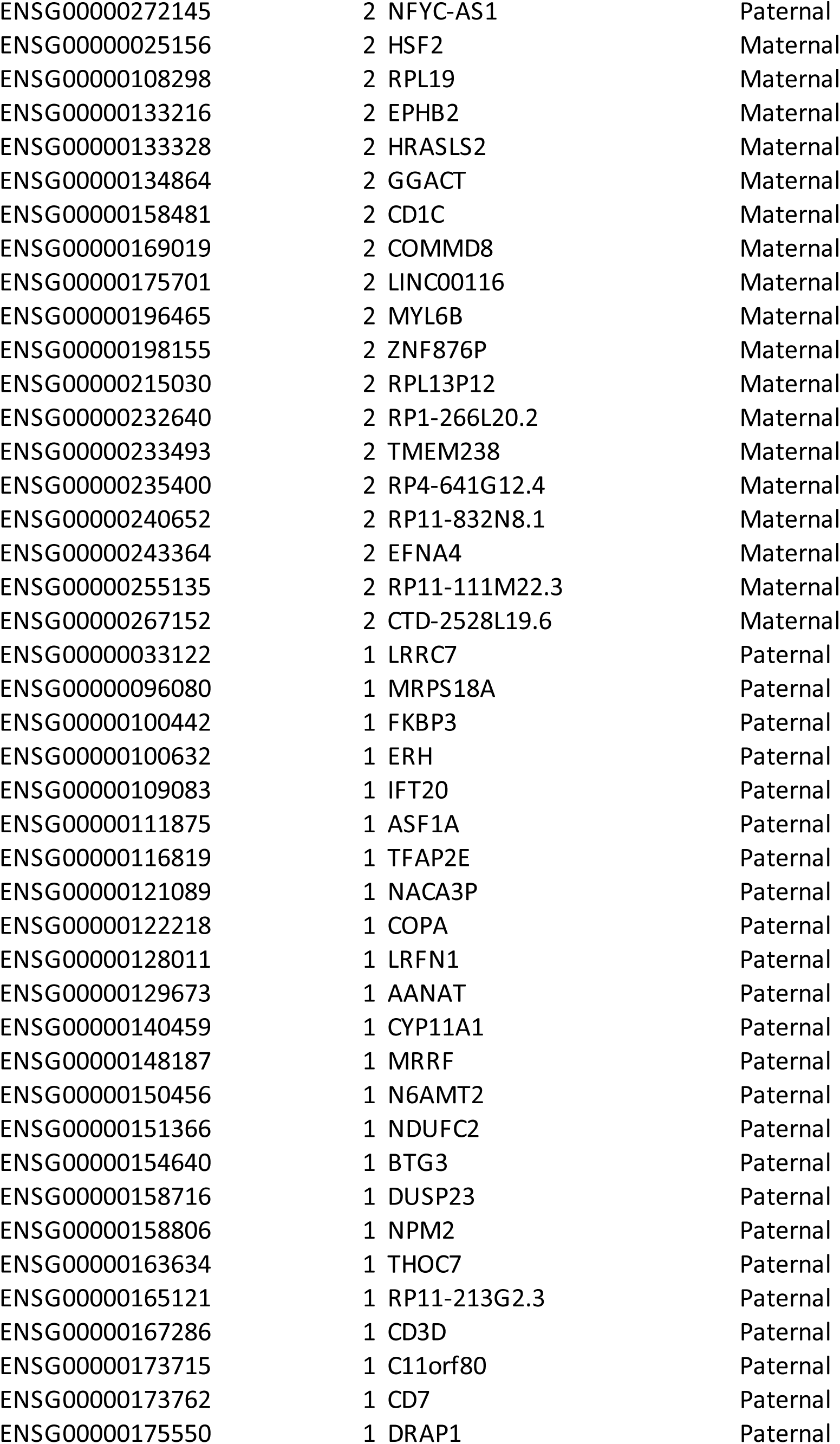
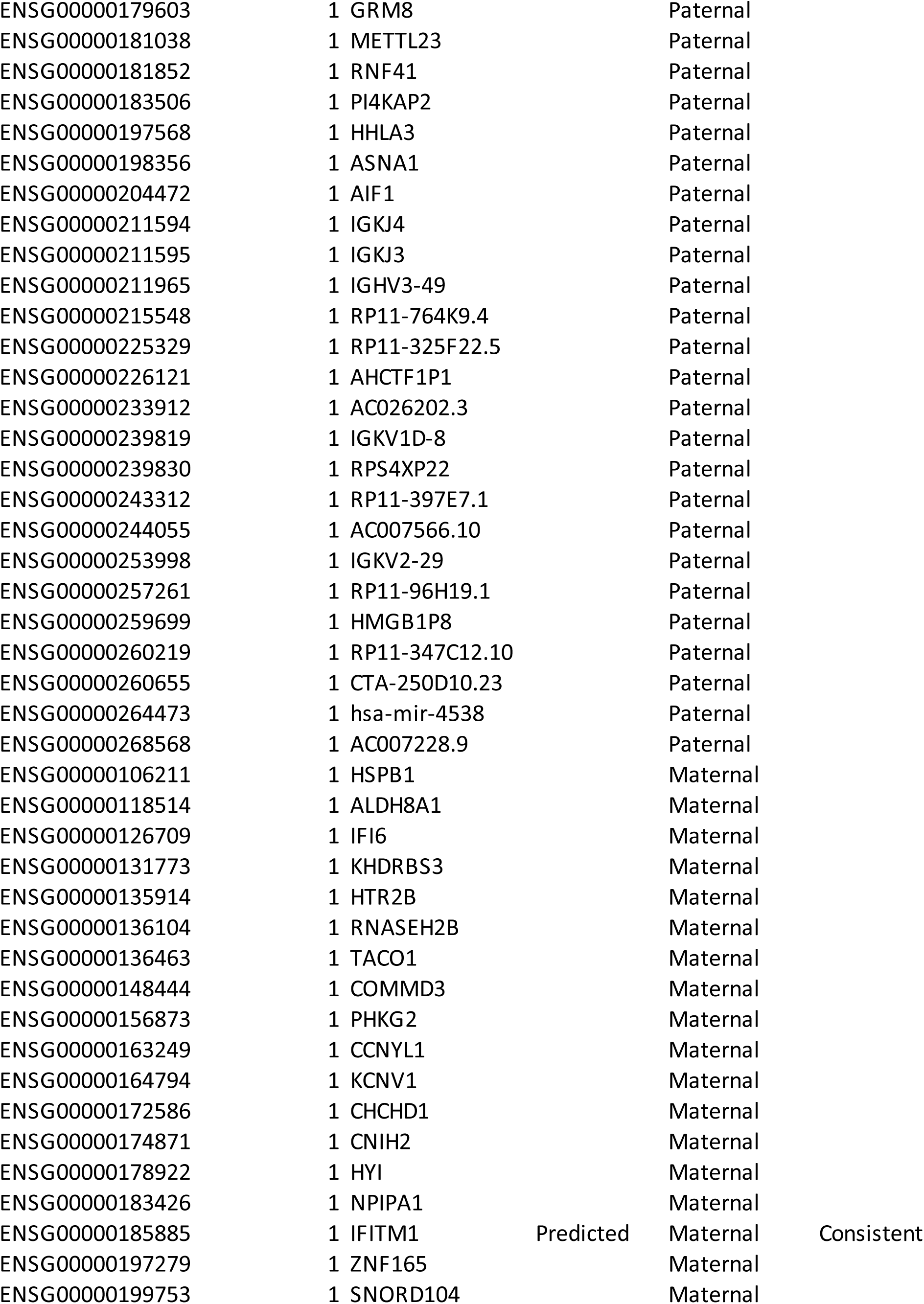
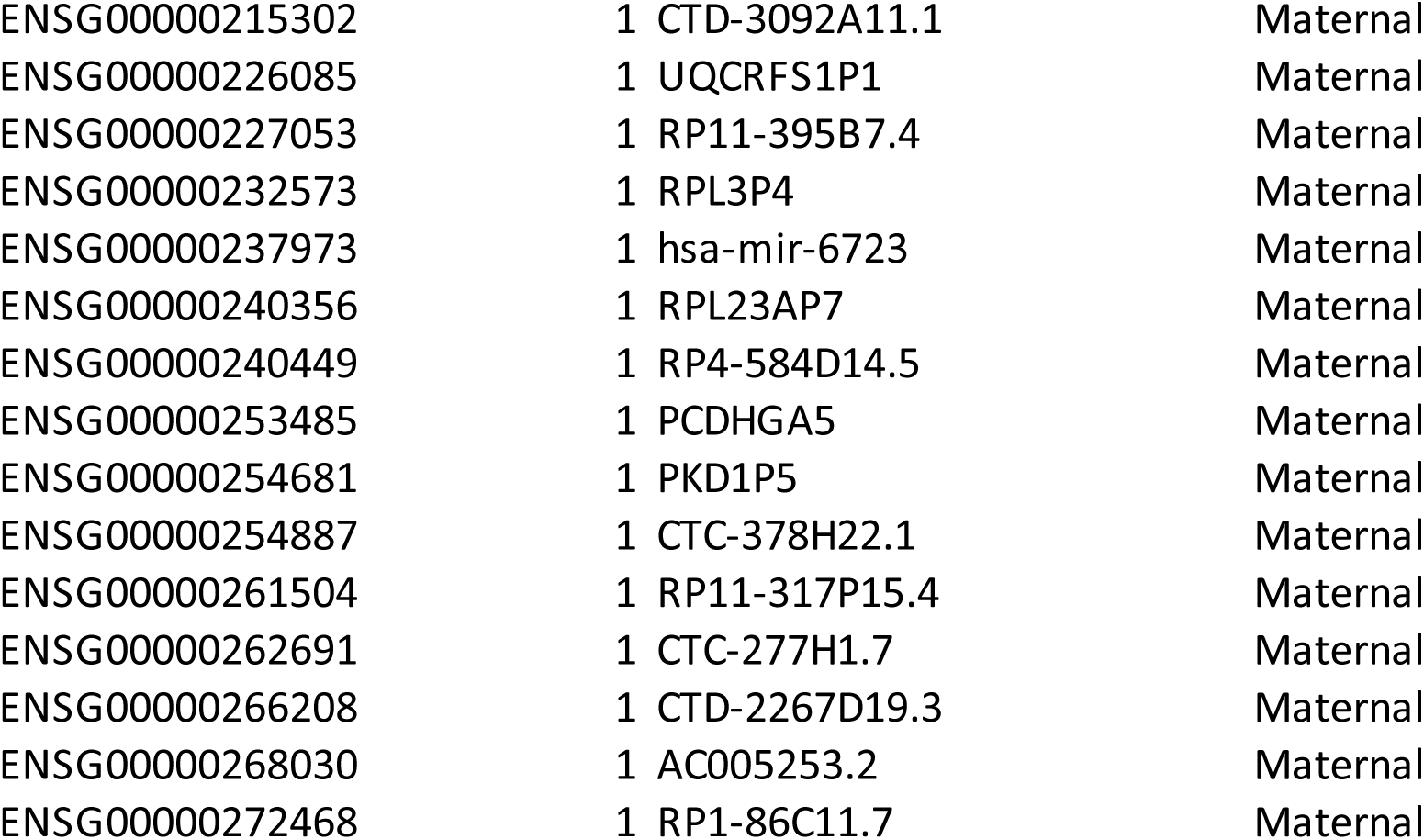
Genes expressed only on maternal or only on paternal haplotypes in LCLs.

